# Rewiring of transcriptional networks as a major event leading to the diversity of asexual multicellularity in fungi

**DOI:** 10.1101/627414

**Authors:** Oier Etxebeste, Ainara Otamendi, Aitor Garzia, Eduardo A. Espeso, Marc S. Cortese

## Abstract

Complex multicellularity (CM) is characterized by the generation of three-dimensional structures that follow a genetically controlled program. CM emerged at least five times in evolution, one of them in fungi. There are two types of CM programs in fungi, leading, respectively, to the formation of sexual or asexual spores. Asexual spores foment the spread of mycoses, as they are the main vehicle for dispersion. In spite of this key dependence, there is great morphological diversity of asexual multicellular structures in fungi. To advance the understanding of the mechanisms that control initiation and progression of asexual CM and how they can lead to such a remarkable morphological diversification, we studied 503 fungal proteomes, representing all phyla and subphyla, and most known classes. Conservation analyses of 33 regulators of asexual development suggest stepwise emergence of transcription factors. While velvet proteins constitute one of the most ancient systems, the central regulator BrlA emerged late in evolution (with the class eurotiomycetes). Some factors, such as MoConX4, seem to be species-specific. These observations suggest that the emergence and evolution of transcriptional regulators rewire transcriptional networks. This process could reach the species level, resulting in a vast diversity of morphologies.

**One-sentence summary:** A study of the evolution of regulators that control the production of asexual spores in fungi.

## Introduction: The importance of asexual development in the fungal life-cycle

According to recent estimates, the fungal kingdom comprises between 2.2 and 3.8 million species, being one of the most diverse clades of eukaryotes (Hawksworth & Lücking 2017). Fungal species have adapted to multiple niches and have developed characteristic cell shapes and morphogenetic/developmental patterns, from single celled yeasts to filamentous multicellular species with a polar mode of cell-growth, or from dimorphic species to flagellated fungi.

Many of these species have been domesticated with some used traditionally for the preservation and/or transformation of foods while others have been optimized to yield valuable biotechnological products (see references within (Johnson 2016; Meyer et al. 2016)). Fungi have also established both symbiotic and antagonistic associations with humans, animals and plants (Hawksworth & Lücking 2017). Over 8,000 fungal plant pathogens are known (Editorial Nature Microbiology 2017) and estimates suggest that fungi (and oomycetes) have an annual impact on major crop yields equivalent to the caloric needs of more than 500 million people (Fisher et al. 2012). Additionally, around 300 fungal human pathogens are known, killing more than 1.5 million people annually. Fungal infections are among the most-rapidly spreading pests (Bebber et al. 2014). This problem is compounded by the simultaneous emergence of the same pathogenic strain in different continents or local episodes of strains from distant locations (Islam et al. 2016; Malaker et al. 2016; Callaway 2016; Lockhart et al. 2017; Bhattacharya 2017).

If a pathogen needs its hosts in order to thrive, theory predicts that the requirement of a minimum host population would cause the pathogen go extinct before the host itself; thus, infection should not be considered a vector that drives extinction (McCallum & Dobson 1995; de Castro & Bolker 2005). Nevertheless, the theoretical models developed by Fisher and colleagues support the idea that fungi pose a greater threat to plant and animal biodiversity than other taxonomic classes of pathogens (Fisher et al. 2012). The authors suggest that this is attributable to some of the biological features of the life-cycle of fungi, such as long-lived environmental stages, broad host ranges in the case of generalist pathogens as well as high virulence and mortality rates. Asexual spores are tightly linked to these traits and this is likely why these propagules have prevailed in evolution as the main vector for the spread of mycoses. Their mass production leads to rapid inter-host transmission rates, which can result in the infection of all individuals before the population is driven to the low densities at which the pathogen can no longer spread (Fisher et al., 2012). Considering the relevance of asexual spores for dispersion and infection, it is important to study how the associated developmental programs have evolved in fungi.

Asexual spores are the last cell-type of a series of genetically programmed morphological transitions leading to the formation, in multiple cases, of complex multicellular structures (see references within (Nagy et al. 2018)). Nevertheless, the morphological diversity of asexual structures generated by the millions of fungal species is vast (Kirk et al. 2008). Fungal asexual spores are produced by four different mechanisms (see references within (Fischer & Kües 2006)): 1) fragmentation of pre-existing hyphal filaments to generate arthrospores or oidia; 2) protoplast contraction and formation of an inner, thickened secondary cell wall within pre-existing vegetative hyphal cells to form chlamydospores; 3) cytoplasmic cleavage within specialized spore mother cells, generating a sporangium and sporangiospores; and 4) localized budding and subsequent constriction from an external sporogenous cell to develop conidiospores or conidia. Furthermore, even if two species develop the same type of asexual spore, they will most probably differ in size, organelle composition and/or morphology. This morphological diversity occurs even among evolutionarily close fungal species, suggesting that fungal asexual development can be considered an example of evolutionary convergence, that is, the generation of structures with different morphologies but with a common function: in this case, the production of spores enabling dispersal to new environments.

With the aim of uncovering the key mechanisms controlling the initiation and progression of asexual multicellularity and hypothesizing how they could contribute to the above-mentioned morphological diversity, we built a database containing the proteomes of 500 fungal species. In parallel, and based on previously published data, we generated a list of 33 proteins known to play key roles in the induction of asexual development, the control of its progression or the balance between asexual and sexual spore production. The conservation of each of these protein sequences was determined for each species in the database, and results were systematically analyzed from an evolutionary perspective. Overall, we propose that emergence of transcription factors (TFs) and network rewiring are major events leading to the diversity of asexual multicellularity in fungi. Results also suggest that asexual development can serve as a model for studying the emergence of CM and the mechanisms controlling it.

## Database generation

The steadily increasing number of sequenced fungal genomes available enables increasingly more informative systematic sequence conservation analyses. These sequencing projects have been generated as a result of individual or collective efforts (Stajich 2017) and have been aggregated and updated in publicly available servers. In this study, the proteomes of 503 fungal species were downloaded from the Emsembl Fungi database (https://fungi.ensembl.org/index.html; release 37; release date, 02/11/2017). These species represented all fungal phyla/subphyla and most classes. The exceptions were neocallimastigomycetes (chytridiomycota); zoopagomycotina and kickxellomycotina (zoopagomycota); coniocybomycetes, lichinomycetes, geoglossomycetes, arthoniomycetes and laboulbeniomycetes (pezizomycotina); archaeorhizomycetes (taphrinomycotina); cryptomycocolacomycetes, cystobasidiomycetes, agaricostilbomycetes, atractiellomycetes, classiculomycetes and tritirachiomycetes (puccioniomycotina); and entorrhizomycetes, and moniliellomycetes (ustilaginomycotina). This meant that, although not in equal number, 65% of the classes included in the tree by Spatafora and colleagues were represented at least by one species in our database (Spatafora et al. 2017). The database contained the proteomes of 344 ascomycota, 114 basidiomycota and 45 early diverging fungal species (1 blastocladiomycota, 4 chytridiomycota, 1 cryptomycota, 24 microsporidia, 14 mucoromycota and 1 zoopagomycota species) (see Table 1). The number of species within each phylum, subphylum, class and order is shown in Table 1 (a supplementary table including the taxonomy of each species of the database, together with the accession number of each proteome, its source, and the first letters of the accession codes used to identify each protein within each proteome is available upon request).

**Table 1:**
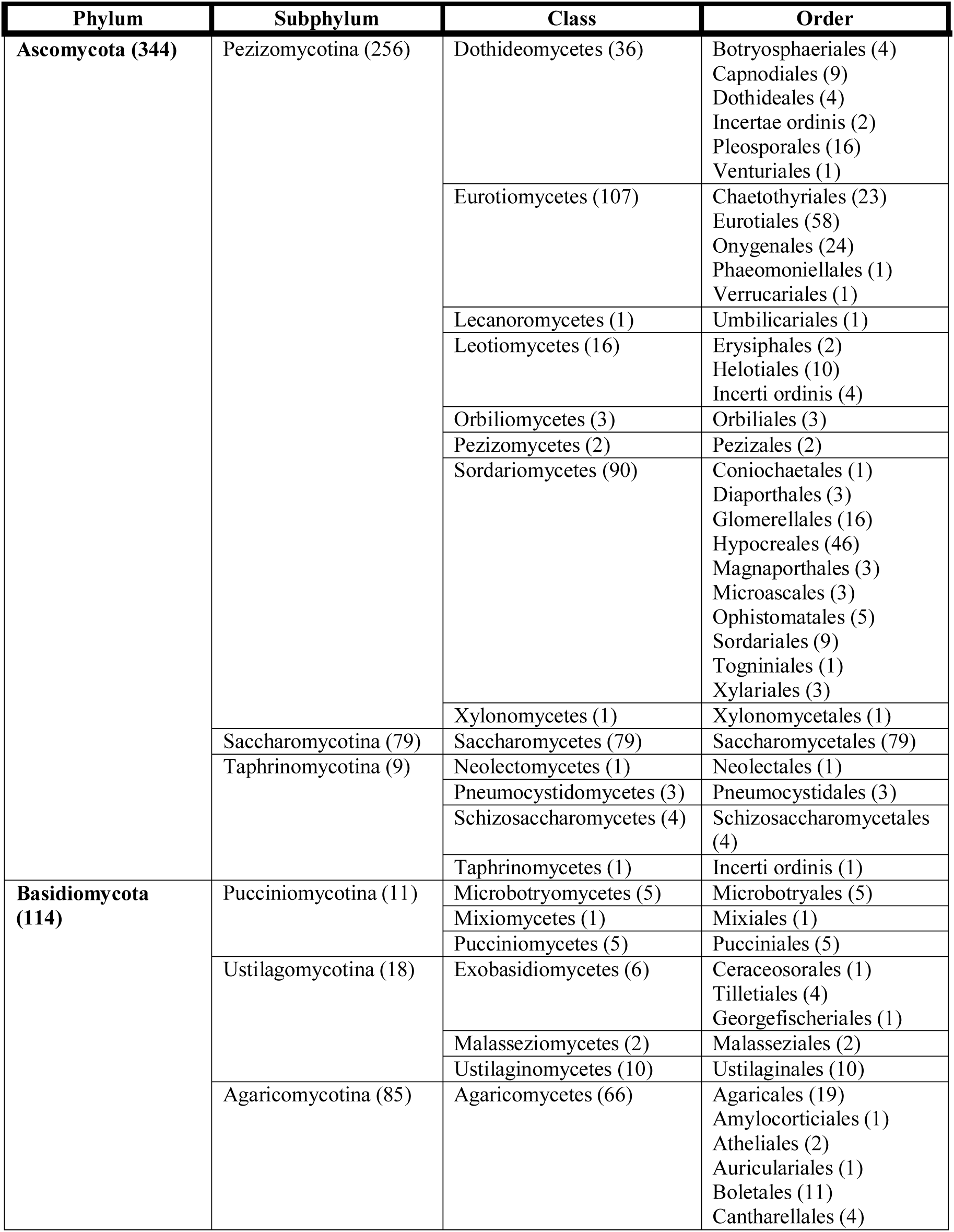
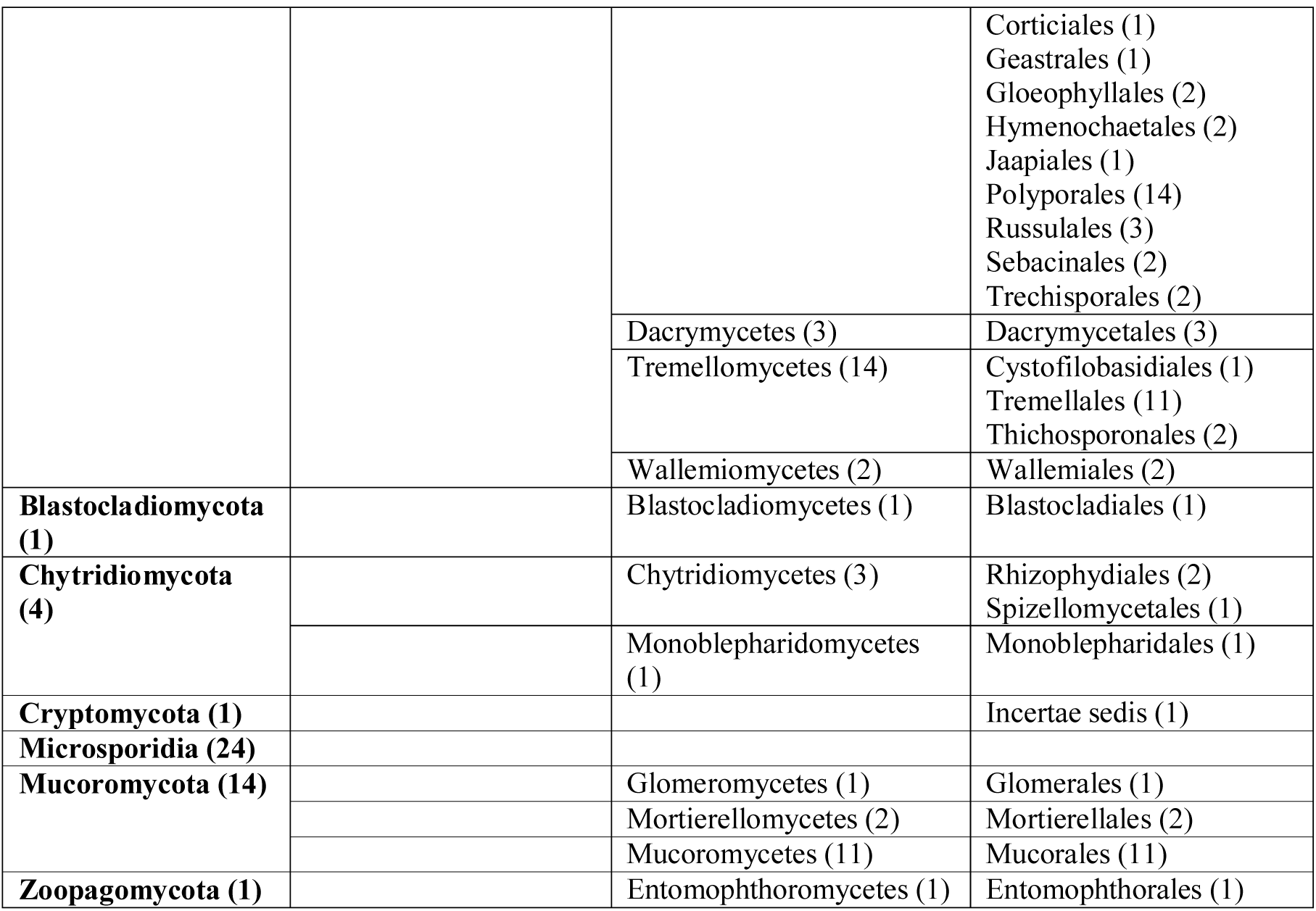
Number of database species (in brackets) in each phylum, subphylum, class and order (503 species). A supplementary table including the taxonomy of each database species, the source of the corresponding proteome and its accession number is available on request.

## Genetic control of asexual multicellularity: Induction, morphological transitions and coordination with the sexual cycle

The literature was reviewed in order to generate a list of regulators of fungal asexual development (Table 2). An enrichment of TFs was noted (28 out of 33; 85 %). These proteins were classified into different groups, according to the role described in the literature (listed in Table 2; displayed in a graphical mode for *Aspergillus nidulans* regulators in Figure 1). First, proteins controlling the balance between sexual and asexual developmental cycles (coded with pink in figures). Second, inducers of asexual development (orange). Third, regulators controlling in space and time the progression of the morphological transitions leading to asexual spore production (green). Finally, those proteins that could not be included in any of the first three groups (yellow). Most of the regulators were originally characterized in *A. nidulans* (eurotiomycetes), which undoubtedly is the most widely used reference organism in the study of asexual CM (Meyer et al. 2016). However, and due to seminal (in the case of *Neurospora crassa*) or more recent (in the case of *Botrytis cinerea* and *Magnaporthe oryzae*) characterization of additional regulators from other species (Selitrennikoff et al. 1974; Cao et al. 2016; Brandhoff et al. 2017), organisms of different fungal clades (Leotiomycetes and Sordariomycetes) have also been included as reference.

**Figure 1:**
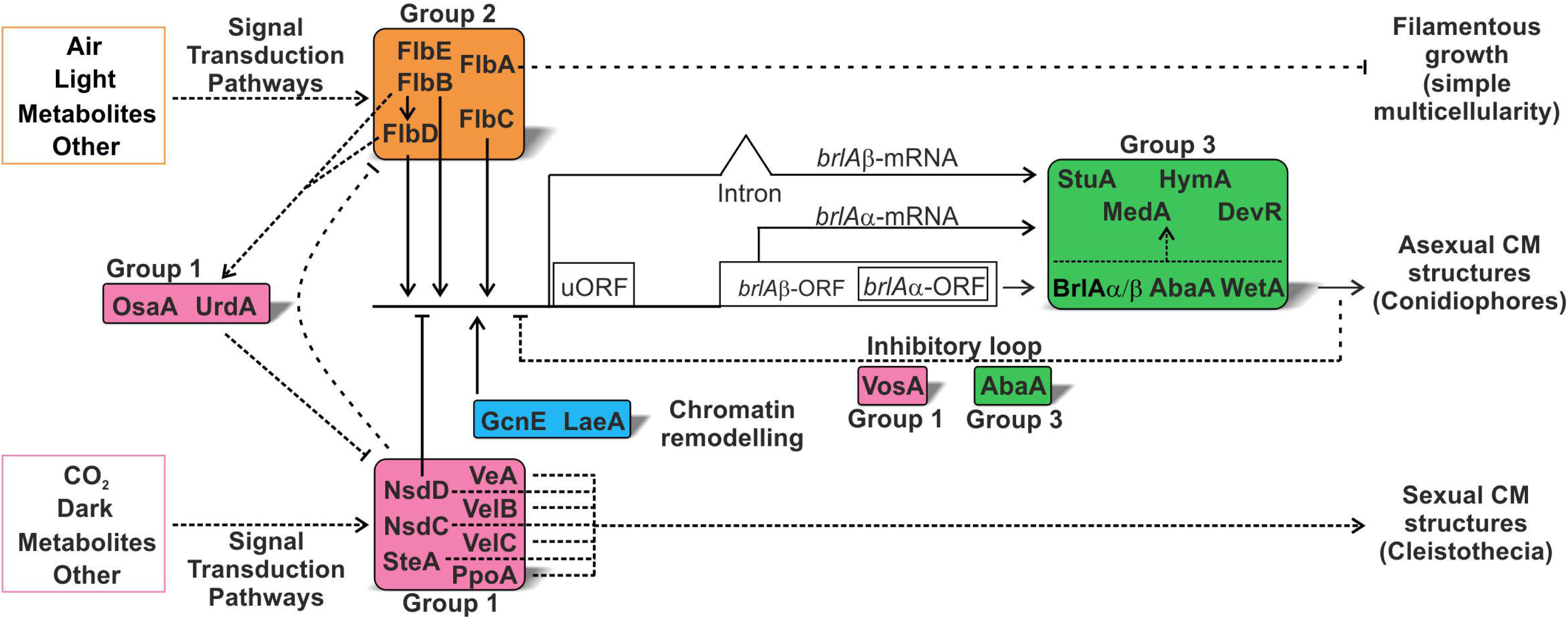
Genetic control of asexual multicellularity in *A. nidulans*. Stimuli such as darkness or carbon dioxide induce the activity of proteins that control the balance between sexual and asexual CM programs (in pink). In general, with the exception of OsaA and UrdA (see below), these proteins inhibit the expression of *brlA* and induce the synthesis of sexual structures known as cleistothecia. Signals such as light and the exposition of vegetative cells to the air environment induce UDA transduction pathways (orange box), resulting in the inhibition of growth (through FlbA) and the induction of *brlA* expression. In addition, UDA activity inhibit sexual development through, at least, UrdA and, probably, also OsaA. Expression of *brlA* is also controlled by chromatin remodelers such as GcnE (and also LaeA in *A. fumigatus*). BrlA activity (first as BrlAβ form then as BrlAα form) initiates the CDP pathway (green box), resulting in the formation of asexual CM structures known as conidiophores. A feedback regulatory loop controlled by VosA informs about the completion of the process by directly inhibiting the expression of *brlA*. The upstream reading frame (uORF) adds a translational layer of control of BrlA activity, since uORFs block ribosomes, decreasing the efficiency of mRNA translation. The position of the arrows indicating binding of each TF at the promoter of *brlA* do not necessarily correspond to the exact position of their binding sites. See references within the main text.

**Table 2:**
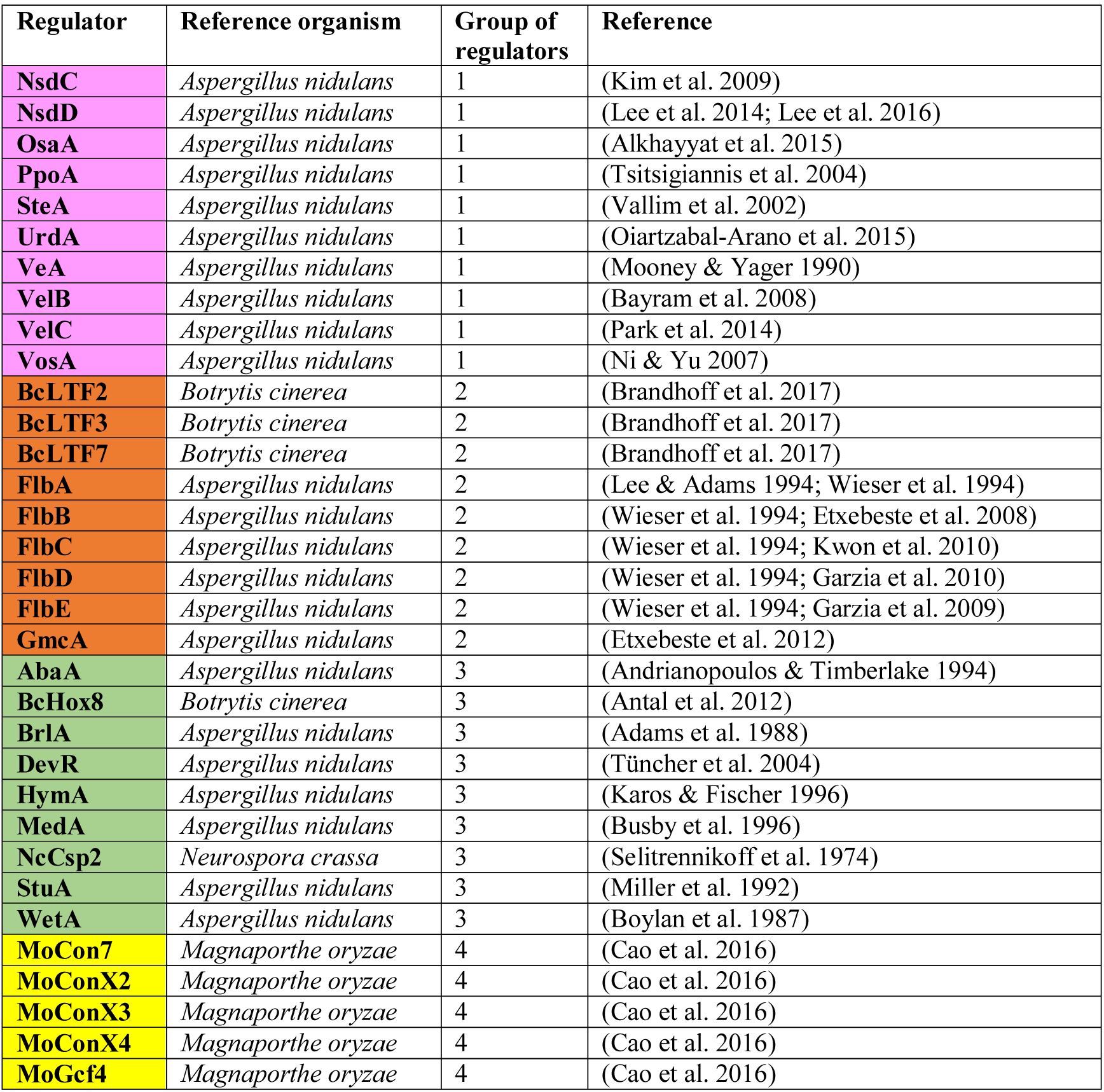
Regulators of asexual development used as query. They are sorted based on their participation in the balance between asexual and sexual developmental cycles (Group 1; pink color will be used), the induction of asexual development (Group 2; orange) and progression of asexual development (Group 3; green). Those proteins not included in any of these three groups are in Group 4 (yellow).

### Group 1: Regulators of the balance between sexual and asexual development

Ten proteins were included in this first group (all characterized in *A. nidulans*; pink color in Figure 1; Table 2). Nine were TFs (VeA, VelB, VelC, VosA, NsdC, NsdD, OsaA, UrdA and SteA). The only non-TF in this group, PpoA, is one of the psi factor-producing oxygenases that control the ratio of sexual to asexual spores (Tsitsigiannis et al. 2004). Among the TFs, VeA, VelB and VelC, together with VosA, belong to the velvet family of transcriptional regulators and are key players in the transduction of light signals as well as the coordination of developmental pathways and secondary metabolism (Ni & Yu 2007; Bayram et al. 2010; Rodriguez-Romero et al. 2010; Ahmed et al. 2013). VeA participates in the transduction of red and blue light signals (Mooney & Yager 1990; Purschwitz et al. 2008) by rapidly varying, in the presence or absence of light, its nuclear/cytoplasmic localization (Stinnett et al. 2006) and its interaction partners (Bayram et al. 2008). One of these partners is VelB, which also interacts with VosA, forming transitory complexes that, as a general rule, promote sexual development and repress asexual development in the dark (see references within (Bayram & Braus 2012)). The nuclear activity of VeA is also controlled by a MAPK module that connects signals received at the tip with the transcriptional control of secondary metabolism (Bayram et al. 2012). This module also controls the induction of sexual multicellularity programs through SteA.

NsdC positively activates sexual development (Kim et al. 2009) while the role of NsdD in the control of multicellularity programs in fungi has been analyzed more deeply (Figure 1; Table 2). Besides being a key activator of sexual development (Han et al. 2001), NsdD directly represses asexual development in *A. nidulans*, by directly binding to the promoter of the master gene *brlA* (see group 3) (Lee et al. 2014; Lee et al. 2016). Finally, WOPR-type and HLH-type TFs OsaA and UrdA were recently characterized functionally in *A. nidulans* (Alkhayyat et al. 2015; Oiartzabal-Arano et al. 2015). The absence of either of them causes very similar phenotypes in which the concentration of asexual spores is significantly reduced while sexual multicellularity is heightened. For this reason, both TFs were classified as repressors of sexual development.

### Group 2: Inducers of asexual development

The regulators included in this second group participate in the transcriptional activation of the genes that control the morphological transitions leading to asexual spore production (group 3; Table 2). In *A. nidulans*, this role is carried out by a group of proteins referred to as UDAs or upstream developmental activators (Wieser et al. 1994; Oiartzabal-Arano et al. 2016). UDAs determine whether the expression of *brlA*, and thus, group 3 genes and asexual spore production, are induced in polarly growing vegetative cells (hyphae) (Figure 1, orange). This is why loss-of-function mutations or deletion of these genes block asexual multicellularity and perpetuate the growth of vegetative cells. Thus, in general, UDA proteins act as signal transducers controlling the transition from simple multicellularity (the mycelium) (Nagy et al. 2018) to complex three-dimensional asexual structures that in the case of *A. nidulans* are formed by the six cell-types that comprise the conidiophore (Adams et al. 1998; Etxebeste et al. 2010).

The most widely characterized group of UDA proteins in *A. nidulans* are the Flb proteins FlbA, FlbB, FlbC, FlbD and FlbE (orange in Figure 1; Table 2) (Wieser et al. 1994). Genetic and molecular analyses placed these regulators into three pathways. FlbA antagonizes the activity of a trimeric G-protein complex formed by FadA, GpgA and SfaD in order to inhibit cell proliferation and induce asexual development (Lee & Adams 1994; Yu et al. 1996). FlbC is a C2H2-type TF that binds the promoter of *brlA*, controlling its expression directly (Kwon et al. 2010). Finally, FlbB, FlbD and FlbE are key players of a characteristic signal transduction pathway connecting the growth region of vegetative cells, the hyphal tip, with nuclei (Oiartzabal-Arano et al. 2016). The bZIP-type TF FlbB is transported to the tip assisted by FlbE (Herrero-Garcia et al. 2015). FlbB migrates from the tip to nuclei, where jointly with the cMyb TF FlbD binds the promoter of *brlA* and induces asexual spore production (Garzia et al. 2010; Herrero-Garcia et al. 2015).

Besides these five regulators, we included in this second group an additional *A. nidulans* protein and three from the leotiomycete pathogen *B. cinerea*. *A. nidulans* GmcA is a predicted glucose-methanol-choline oxidoreductase required for asexual multicellularity under alkaline pH conditions (Etxebeste et al. 2012). BcLTF2 and BcLTF3 are recently characterized, inter-dependent, light-induced TFs from *B. cinerea* (Brandhoff et al. 2017) while BcLTF7 is the ortholog of MoConX7, a C2H2 zinc-finger TF whose absence reduces conidia production in *Magnaporthe oryzae* by more than 60% (Cao et al. 2016).

### Group 3: Spatiotemporal control of the synthesis of asexual multicellular structures

From the above-referenced literature, it can be concluded that, in *A. nidulans*, a first layer of transcriptional control at the promoter of the master gene *brlA* is exerted by both activators and repressor TFs belonging to different transduction pathways (Lee et al. 2016). Once *brlA* expression is induced, a cascade of transcriptional regulators is activated (green in Figure 1; Table 2) (Adams et al. 1998). This pathway is known as CDP or Central Developmental Pathway (Mah & Yu 2006) and controls the expression of an array of genes required, for example, in the synthesis of the cell-wall or pigmentation of spores (see references within (Adams et al. 1998); these auxiliary proteins have not been considered in the analyses below).

There are additional layers of transcriptional control of the expression of *brlA*. The second one is the existence of two transcripts of *brlA*, *brlA*α and *brlA*β (Han et al. 1993). *brlA*β codes for a polypeptide 23 amino acids longer than BrlAα. Each isoform is required at different stages of asexual multicellular development: *brlA*β mutants are blocked at early-mid asexual CM development while *brlA*α mutants fail to proceed with the production of asexual spores (Fischer & Kües 2006). Transcription of *brlA*β starts upstream to that of *brlA*α, and the transcript contains an intron in the 5’-UTR region. Transcription of *brlA*α starts within the intron of *brlA*β (our RNAseq results; (Garzia et al. 2013; Oiartzabal-Arano et al. 2015)).

A third layer is established by GcnE, the histone acetyltransferase subunit of the Spt-Ada-Gcn acetyltransferase (SAGA) complex, which strongly suggests that histone acetylation and chromatin remodeling at the promoter of *brlA* are key events in the development of asexual CM structures (Cánovas et al. 2014). This is supported by a recent study in *A. fumigatus*, which reports that LaeA prevents heterochromatic marks in the promoter of *brlA*, allowing its activation (Lind et al. 2018).

Fourthly, feedback regulatory loops on *brlA* mediated by group 1 (VosA) or group 3 (AbaA; see below) TFs have also been described, refining the expression and activity of BrlA isoforms and informing about the completion of the process (Aguirre 1993; Ni & Yu 2007). Finally, the presence of a uORF just 3’ of the initiation site of *brlA*β transcript adds a translational layer of control of BrlA concentration in the cell.

It remains to be determined whether BrlA counterparts and associated complex regulatory mechanisms are coded in the genomes of species outside the order eurotiales (see next sections). However, *brlA* constitutes a control point in *A. nidulans* and, as far as known, it is the only way to induce the CDP pathway and asexual CM, thus ensuring that all the cell-types that form the conidiophore are generated correctly in space and time. The conservation of TFs AbaA and WetA, which together with BrlA form the backbone of the CDP pathway, and StuA and MedA, has also been analyzed (see also (Ojeda-López et al. 2018)). AbaA is a TF of the ATTS family (Andrianopoulos 1991) required in the terminal stages of asexual development (Andrianopoulos & Timberlake 1994) while WetA controls gene expression in asexual spores and is required for their integrity and the deposition of diverse metabolites (Sewall et al. 1990; Marshall & Timberlake 1991; Wu et al. 2018). StuA and MedA purportedly control the activity of CDP factors in space and time (Miller et al. 1992; Busby et al. 1996).

Four additional proteins were included in this third group: the bHLH-type TF DevR, the homeobox TF BcHox8, HymA and the CP2-like TF NcCsp2. The deletion mutant of *devR* initiated conidiophore development in *A. nidulans* but failed to proceed with conidia production and instead generated secondary conidiophores (Tüncher et al. 2004). The absence of *B. cinerea* BcHox8 caused the generation of conidiophores and conidia with abnormal morphology (Antal et al. 2012). HymA has been included in this group as a protein that, being conserved in higher eukaryotes, is necessary at middle-late stages of asexual CM development (Karos & Fischer 1996). *N. crassa* has been an important model for the study of asexual development in fungi and multiple regulators of this process are known (see references within (Park & Yu 2012)). Among them, NcAcon3, required for the formation of major constriction sites, is the ortholog of MedA while NcCsp1, which is required for the separation of conidia, is the ortholog of BcLTF3 (see Group 2). In this study, we used NcCsp2 as a query, a TF also required for the separation of conidia (Selitrennikoff et al. 1974).

### Group 4: Other regulators

Some of the regulators of asexual development analyzed in this review do not fit in any of the previously defined groups mainly because their role has been characterized in model fungi distant from *A. nidulans*. Cao and collaborators carried out a comprehensive characterization of the roles of 47 C2H2-type TFs in the development and pathogenicity of *M. oryzae* (Cao et al. 2016) and described that individual deletions of several of them caused at least an 80% decrease in conidia production without causing a significant inhibition of colony growth. Therefore, MoGcf4, MoConX2, MoConX3, MoConX4 and MoCon7 were included in the list of TFs analyzed in this work.

## TFs controlling multicellular development in fungi emerged gradually in evolution

The sequences of the 33 regulators described above (Table 2) were retrieved from FungiDB (http://fungidb.org/fungidb/), NCBI (https://www.ncbi.nlm.nih.gov/) or AspGD (http://www.aspgd.org/) databases. The 503 fungal proteomes were divided into sub-databases prior to analysis: 1) agaricomycotina (85 species); 2) blastocladiomycota (1), chytridiomycota (4), cryptomycota (1) and zoopagomycota (1); 3) microsporidia (24); 4) mucoromycota (14); 5) pezizomycotina (256); 6) puccioniomycotina (11) and ustilagomycotina (18); 7) saccharomycotina (79); and 8) taphrinomycotina (9). The sequence of each regulator was individually processed as the query in BLAST searches against each sub-database. Additional functional annotation was performed by searching subsets of the EnsemblFungi database (https://fungi.ensembl.org/index.html) using emapper-1.0.3 (Huerta-Cepas et al. 2017). Additionally, HMM profiles for regulators were obtained using sequence searches of the eggNOG 5.0 database (http://eggnogdb.embl.de/#/app.home) (Huerta-Cepas et al. 2016; Huerta-Cepas et al. 2019).

All hits from each query were analyzed based on the following criteria. First, the presence, located in a similar position within the hit, of the characteristic domain(s) of the query. Second, a threshold coverage value was established and used as criteria for exclusion. In general, every hit with a query coverage below 35% was not considered as homologous and was consequently excluded from the analysis. Exceptions to this rule were included. A minimum coverage of 30% was established for FlbD. For MoConX3 a minimum coverage was not considered, prioritizing the conservation of its two transcriptional regulatory domains at N- and C-termini of the protein. For SteA, the conservation of its two transcriptional regulatory domains at N- and C-termini or a minimum coverage of 35% was accepted. Third, BLASTing each specific hit against the genome database of the query species (www.aspgd.org or www.fungidb.org) had to return the query as the first hit (confirmatory reverse retrieval).

The application of our criteria, especially the confirmatory reverse retrieval, led us to conclude, for example, that VeA is conserved, almost exclusively, in pezizomycotina, which contrasts with previous studies that described its presence in species outside of this subphylum (Bayram & Braus 2012). Interestingly, we obtained the same hits for VeA and VosA in most species outside of pezizomycotina. In the case of VelC, the presence of the velvet domain in the C-terminus of the protein was prioritized over the confirmatory reverse retrieval results (velvet domains of both VeA and VosA are located in the N-terminus).

Results strongly suggest that three proteins belonging to the velvet family of regulators (VelB, VelC and VosA) are present in nearly all fungal clades, indicating that this is one of the most ancient fungal systems coupling signal transduction with developmental and metabolic control. Ojeda-López and colleagues analyzed the conservation of the four velvet proteins of *A. nidulans* in 54 proteomes representing different fungal phyla, subphyla and classes, and proposed different hypotheses on the evolution of this family of regulators (Ojeda-López et al. 2018).

By applying these criteria, we generated lists of orthologs (supplementary table available on request), which include the accession code of each hit, coverage, expected and score values, as well as the best hit resulting from a BLAST analysis of that sequence against the reference database (the lowest expected value was 3,00e^-05^, corresponding to the *Thielaviopsis punctulata* ortholog of FlbE KKA29905). The heatmap in Figure 2 was generated with the expect values of all hits of those queries considered as orthologs (supplementary table available on request).

**Figure 2:**
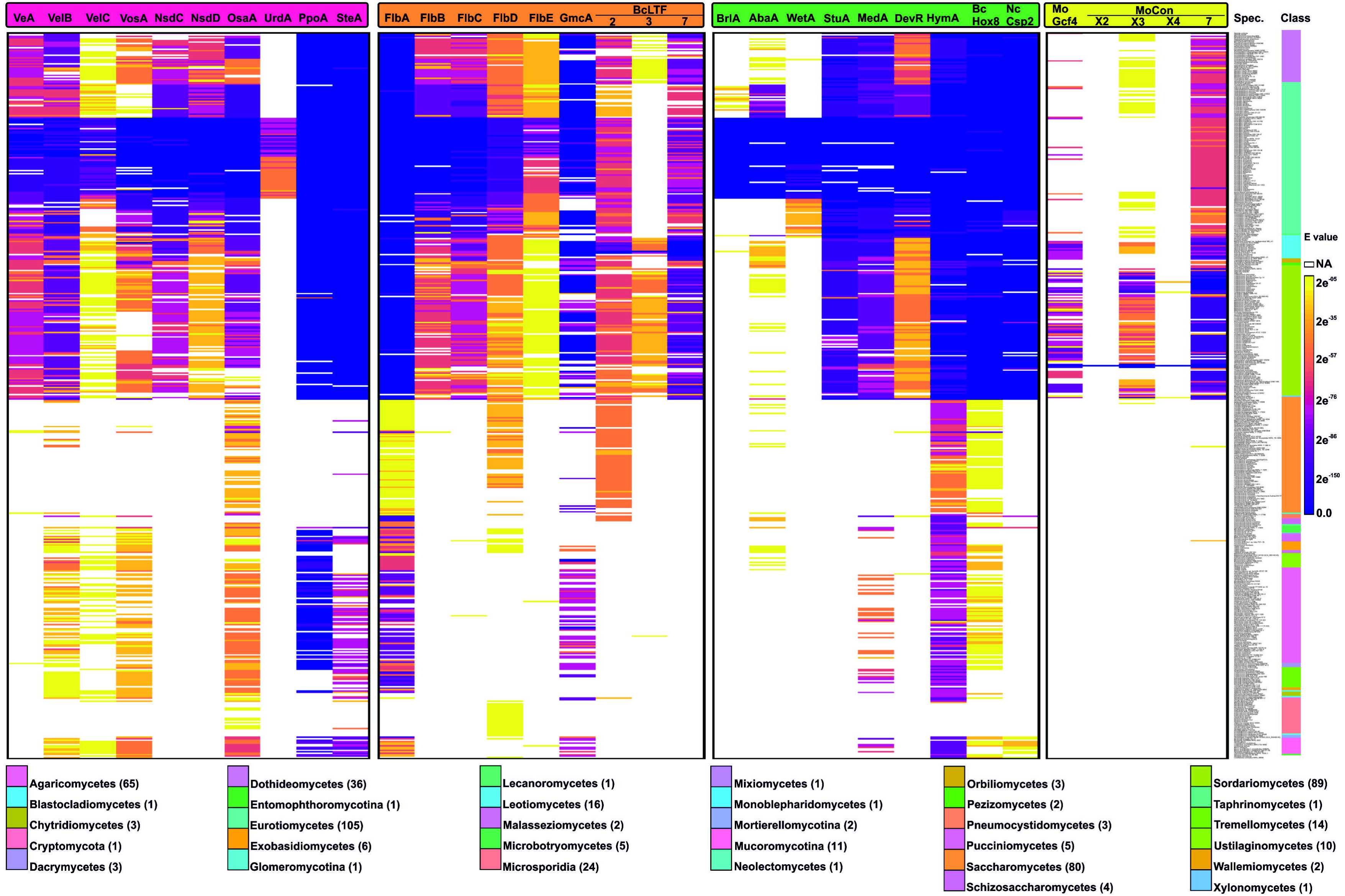
Conservation of developmental regulators in fungi. Heatmap showing the expected (e) values of the orthologs of each developmental regulator identified in each of the species of the database. White color (NA) indicates an absence of an ortholog in that species or that a hit was discarded because it was below our threshold criteria (see main text). The color bar on the right indicates the class (labeled at bottom of figure) of those species.

The heatmap in Figure 3 describes the fraction of species within each class with an ortholog of each developmental regulator (the corresponding table is available on request). Overall, three patterns of conservation applicable to the four groups of regulators described were observed. First, the conservation of 13 of the regulators (all but one being TFs) was limited to all or nearly all classes within ascomycota or pezizomycotina. Second, six TFs were conserved only in specific classes within pezizomycotina. Here we observed sub-patterns of conservation, with some of these regulators being conserved only in one class or specific orders, families, or even species (see Figure 2), while others were found in more than one. Finally, we found that 14 of the proteins analyzed were also conserved in orders outside of ascomycota/pezizomycotina clades, suggesting a more ancient emergence in evolution (see below).

**Figure 3:**
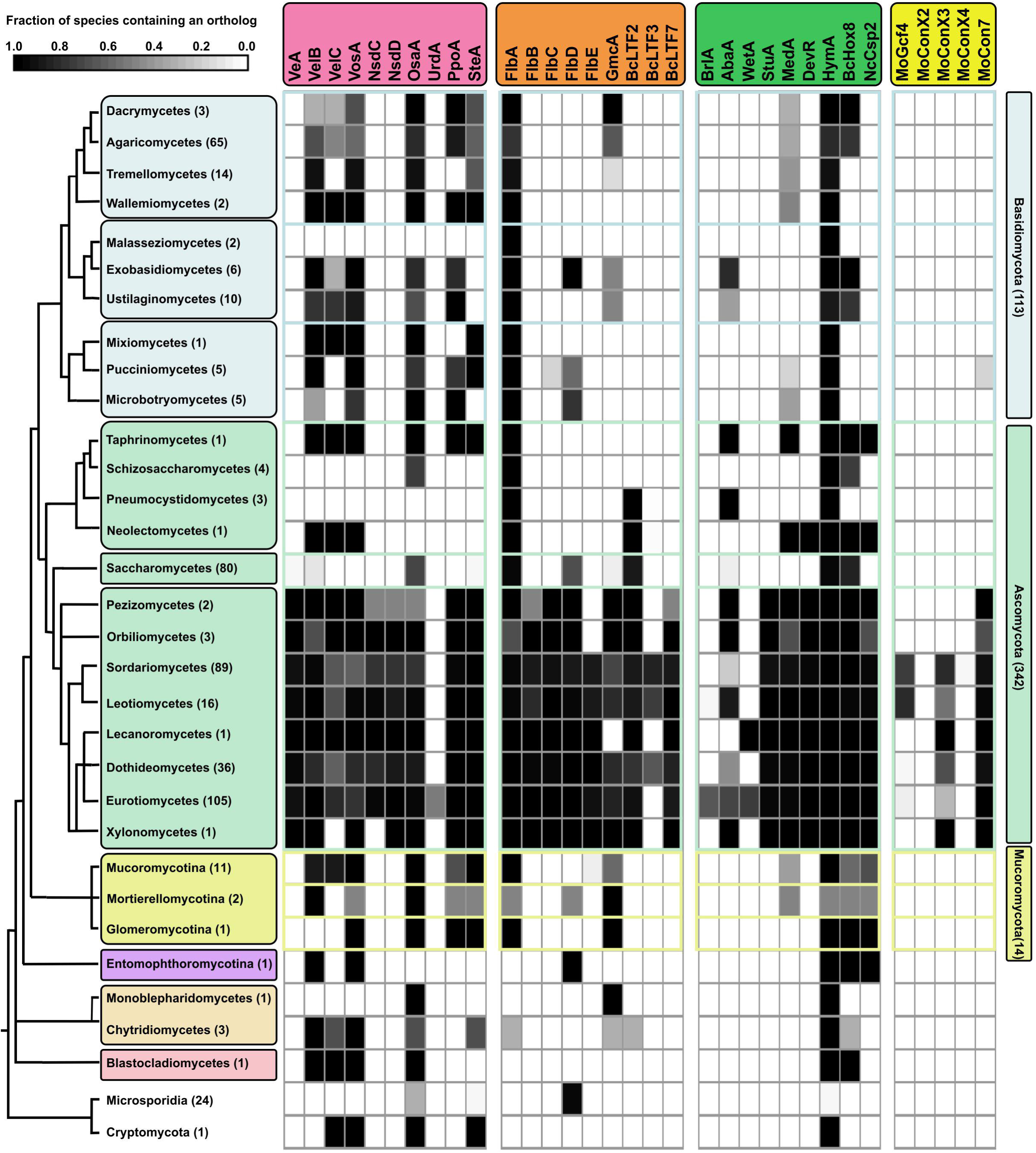
Conservation of regulators of asexual development in those classes represented in our database. Blue color on the phylogenetic tree represents the three subphyla within basidiomycota (agaricomycotina, ustilaginomycotina and pucciniomycotina, respectively); green represents the three subphyla within ascomycota (taphrinomycotina, saccharomycotina and pezizomycotina, respectively); yellow indicates mucoromycota; purple, zoopagomycota; beige, chytridiomycota; and pink, blastocladiomycota. The tree was manually drawn based on that published by (Spatafora et al. 2017). The number of representative species of each class in our database is indicated in brackets. The greyscale indicates the fraction of species within each class containing an ortholog of each developmental regulator (see also Table S1). Conservation was determined by BLAST and eggNOG analyses, according to the criteria described in the main text.

By identifying the most phylogenetically ancient classes in which orthologs of the analyzed query proteins could be detected, we tried to estimate the relative phylogenetic age (phylostrata) or point in which each of the regulators could have emerged in evolution (Krizsán et al. 2019). With this aim, the conservation patterns shown in Figure 3 were placed on a phylogenetic tree based on Spatafora and colleagues as the reference (Figure 4) (Spatafora et al. 2017). A protein was considered as conserved in a specific class when orthologss were identified in at least 40% of the total number of species in our database representing that class (Nguyen et al. 2017).

**Figure 4:**
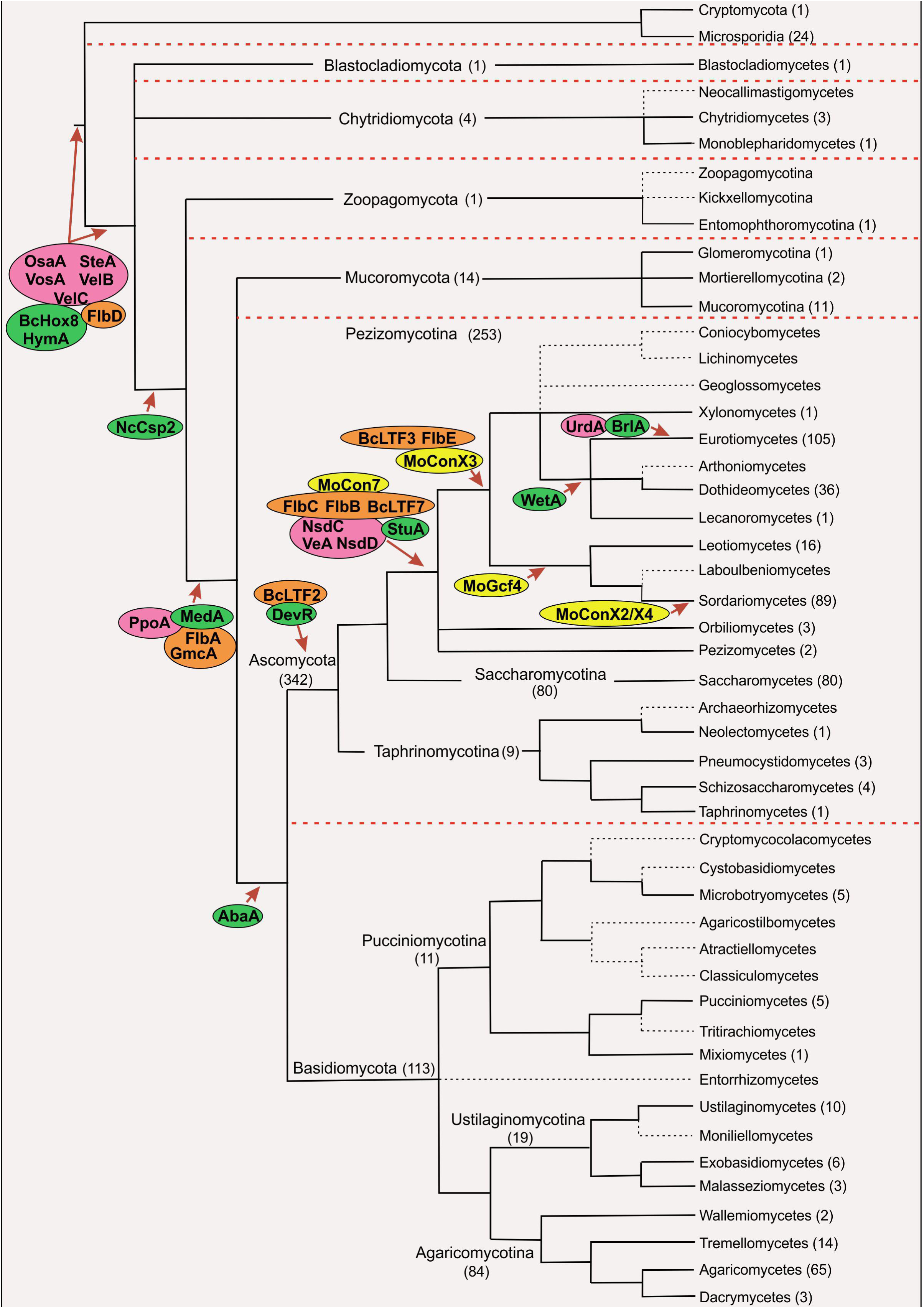
Emergence of regulators of asexual development. By drawing a phylogenetic tree based on Spatafora and colleagues as a reference (Spatafora et al. 2017), the figure shows the number of species (in brackets; a total of 500) of each phyla, subphyla and class represented in the database of fungal proteomes used in this study. Classes not represented in our database are indicated by black dotted lines. The red dotted lines have been included to separate major clades. The locations of the regulators of asexual development in the tree are based on the results obtained in BLAST and eggNOG analyses and suggest an approximate period for their emergence. A protein was considered as conserved in a specific class when orthologs were identified in at least 40% of the total number of species representing that class in our database. See also Figures 2 and 3.

Although in some cases it is difficult to track the emergence of specific regulators due to the under-representation of specific classes in our database, Figure 4 strongly suggests that TFs controlling complex multicellularity in fungi emerged sequentially together with the split of major clades. Three of the velvet proteins of *A. nidulans* (VosA, VelB and VelC), together with additional TFs such as FlbD, OsaA and BcHox8, as well as HymA, are apparently the most ancient regulators of fungal development (a BLAST of HymA sequence against non-fungal UniProtKB taxonomic subsets at https://www.ebi.ac.uk/Tools/sss/ncbiblast/ confirmed that it is conserved in higher eukaryotes as well (Karos & Fischer 1996)).

The emergence of multiple regulators in the ascomycota or pezizomycotina clades is in line with the observation by Krizsán and coworkers that there is a characteristic enrichment of agaricomycetes-specific TFs (Krizsán et al. 2019) and suggests that the development of CM patterns in these phyla and subphyla was accompanied by the emergence of several new developmental regulators enabled by this TF expansion (Figure 4). Also in line with the hypotheses of Nguyen and collaborators based on the genome sequencing of *Neolecta irregularis* (a taphrinomycete which develops sexual CM structures), regulators such as BcLTF2 or DevR probably are ancestral in the ascomycota, and DevR (but not BcLTF2) have probably been lost during simplification of budding and fission yeasts (Nguyen et al. 2017). As far as we know, there is no report on any role in sexual (sclerotial) development for the regulator of *B. cinerea* conidiation BcLTF2 (Brandhoff et al. 2017). However, BcLTF3 has been described as a TF with a dual function in development (see below). Of note is the conservation pattern of NcCsp2, since it is conserved in pezizomycotina, mucoromycota, zoopagomycota and the taphrinomycotina *Neolecta irregularis* and *Saitoella complicata*, but not in saccharomycotina, the rest of taphrinomycotina or basidiomycota species analyzed in this work. This suggests an early emergence of NcCsp2 and the progressive loss in basidiomycota, saccharomycotina and specific clades within taphrinomycotina.

MoConX3, FlbE and BcLTF3 (the latter has a dual function in the control of development since it is essential for conidiogenesis but represses conidiophore development by repressing *Bcltf2* in light and darkness; see (Brandhoff et al. 2017)) may have emerged after the split of orbiliomycetes and pezizomycetes, while *M. oryzae* MoGcf4 is conserved in leotiomycetes and sordariomycetes. Furthermore, *A. nidulans* UrdA and BrlA, as well as *M. oryzae* MoConX2 and MoConX4 seem to be specific to eurotiomycetes and sordariomycetes, respectively. However, there is a variable degree of conservation among these regulators. UrdA is conserved exclusively in eurotiales, showing a variable degree of conservation in this order (see Figure 2). The C2H2-type regulator MoConX2 shows a high sequence divergence compared to the best hits in other sordariomycetes, being located in an independent clade (see the analysis of MoConX2 sequence in Figure S1), while we found only three putative ortholog sequences for MoConX4, all of them with low coverage and expected values (see Figure 2). This observation strongly suggests that the emergence of specific TFs controlling asexual multicellularity may be recent and may lead to further morphological diversification and specialization. The possibility of these TFs emerging sooner in evolution and being gradually lost in almost all species is unlikely.

Presence of BrlA orthologs is detected in the order eurotiales within eurotiomycetes (also probably in some chaetothyriales species and the single verrucariales species analyzed, *Endocarpon pusillum*), but not in onygenales (de Vries et al. 2017). The fact that BrlA emerged in evolution much later than TFs that in *A. nidulans* are located either upstream (FlbA-E) or downstream (AbaA, WetA, StuA, MedA) in the genetic pathways inducing/controlling the generation of asexual CM structures suggests that its emergence had a profound effect in the modification of pre-existing democracy/hierarchy networks (see Discussion).

## Discussion

Multicellularity represents a key landmark in the evolution of organisms. Complex multicellularity (CM) is differentiated from simple multicellularity based on features such as the proportion of cells in direct contact with the environment, cell adhesion and communication, the existence of a developmental program controlling morphological transitions, programmed cell death events or the generation of 3D structures (see references within (Nagy et al., 2018)). CM has emerged at least five times in evolution, one of them in fungi. There are two types of CM programs in fungi, sexual and asexual cycles. However, investigations into the fundamentals of the emergence and evolution of CM in fungi has been dominated by studies of sexual development. Here we propose that the study of the emergence/evolution of regulators that control the initiation/progression of asexual development in specific fungal lineages can be an informative way to elucidate the mechanisms enabling asexual CM and potentially leading to evolutionary convergence. For example, the generation of conidiophores meets most of the traits of CM mentioned above. Although all cell-types are in direct contact with the environment, which contrasts with (asexual or sexual) fruiting bodies, conidiophores are 3D structures generated following a pre-defined genetic program that is tightly connected to programmed cell death pathways (Mims et al. 1988; Adams et al. 1998; Pócsi et al. 2009; Gonçalves et al. 2017). Two additional advantages reinforce the usefulness of asexual development as a model for the study of CM. One is the limited number of cell-types involved. In *A. nidulans* conidiophores, the foot-cell (the base of the structure), the stalk (which grows polarly from the foot-cell) and the vesicle (the result of the isotropic growth of the tip of the stalk) form a single unit (Mims et al. 1988). A multipolar budding process at the vesicle generates metulae, which subsequently bud into two phialides, each one generating long chains of asexual spores (conidia). Secondly, and again using the example of *A. nidulans* conidiophores, it can be suggested that the timely generation of each cell-type is controlled by a limited number of transcriptional networks and TFs (Etxebeste et al. 2010).

Most of these TFs (a significant number at least) are known in *A. nidulans* and the functional characterization of their roles in asexual development has enabled their assignment to pathways and the elucidation of their hierarchical or democratic relationships (Bar-Yam et al. 2009; Jothi et al. 2009). For example, *brlA* expression is dependent on the activity of UDA TFs (hierarchy) while, the expression, protein levels, localization and transcriptional activity of both FlbB and FlbD depend on the activity of each other (democracy) (our unpublished results; (Garzia et al. 2010)). Our results strongly suggest that UDA TFs and others controlling the initiation, progression or balance of asexual development are widely conserved in most classes within pezizomycotina. Nevertheless, the functional characterization of several of these orthologs clearly shows that sequence conservation does not guarantee the conservation of the same role in the control of development. For example, the deletion of the orthologs of *flbB*, *flbE* and *fluG* did not alter conidia production or morphology in *Fusarium graminearum* while deletion of *fgflbC* caused approximately a 50% decrease in conidia production and deletion of *fgflbD* completely blocked the induction of asexual development (Son et al. 2014). Thus, a democratic relationship between FgFlbB and FgFlbD in the control of conidiation in this sordariomycete is very unlikely. Tiley *et al*. have recently shown that the deletion of the *flbB* ortholog in the dothideomycete *Zymoseptoria tritici* had no effect on asexual spore production or pathogenicity (Tiley et al. 2018). Furthermore, deletion of *M. oryzae MoflbB* exacerbated conidiation instead of inhibiting it (Tang et al. 2014), suggesting that MoFlbB performs an opposite role compared to the *A. nidulans* ortholog. FlbB orthologs conserve different domains and residues compared to related TFs that control the response to oxidative stress (Vivancos et al. 2004; Cortese et al. 2011; Herrero-Garcia et al. 2015). This may indicate that FlbB orthologs retain mechanistic features enabling them to control signal transduction, such as importation into nuclei or transcriptional regulation, but can participate in the alternate cellular process in different species. Additionally, the emergence of BrlA in eurotiales (and apparently some chaetothyriales) may have re-directed the role of FlbB towards the induction of asexual development (see below).

Similarly, deletion of the *N. crassa* ortholog of *flbC*, *flb-3*, has pleiotrophic effects, affecting hyphal morphology as well as sexual and asexual development (Boni et al. 2018). As shown in Figure 1, FlbC regulates *brlA* expression in *A. nidulans* by direct binding to its promoter (Kwon et al. 2010). However, as a sordariomycete, *N. crassa* lacks a *brlA* ortholog and Flb-3 has developed the ability to bind the promoters of *aba-1*, *wet-1* and *vos-1*, which code for proteins orthologous to *A. nidulans* CDP TFs AbaA, WetA and VosA. Unexpectedly, the authors reported that these three genes are apparently not required for development in *N. crassa* (Boni et al. 2018). As evidence of the complexity in the organization of the networks controlling development in fungi, in the insect pathogen *Beauveria bassiana*, a sordariomycete that consequently lacks a *brlA* ortholog, deletion of *BbwetA* and *BbvosA* affects conidia number and quality (Li et al. 2015). In a context in which sequence conservation does not correlate with conservation of the same cellular function, and new TFs controlling development emerge as the different fungal orders, families, genera and probably even species themselves differentiate implies that broadening the search for mutants impaired in asexual development is necessary in as many reference systems as possible. Thus, mutant screenings such as the one carried out in *B. cinerea* for the identification of light induced TFs (Cohrs et al. 2016; Schumacher 2017; Brandhoff et al. 2017) or the identification and characterization of proteins controlling asexual sporulation in agaricomycetes (reviewed by (Kües et al. 2016)) serve as examples and must be acknowledged. Overall, these examples suggest that the application of known activities and relationships of developmental regulators from one model to related species cannot be treated as a given, and may misdirect research. Furthermore, collaborative initiatives should integrate the data generated not only from evolutionary and network organization perspectives, but also with the understanding that there is the potential of discovering novel therapeutic targets within those pathways, since, after all, they control the spread of mycoses. In this context, and considering the fact that asexual reproduction is prolific and results in wide dispersion, random mutation of spores could promote speciation.

Recent reports propose that an increase in the number of protein-coding genes is not the only cause of the development of CM in fungi, indicating that there are other sources of genetic innovations (Nagy 2017; Nguyen et al. 2017). The observations described in this text lead us to propose that asexual CM in pezizomycetes evolved convergently, at least, as the result of a random and stepwise emergence of TFs and additional regulators. The emergence or re-adaption of regulators within the pre-existing transcriptional networks would cause their rewiring, meaning that the great diversity of asexual morphologies in fungi is the result of random re-organizations of genetic pathways. In fact, inefficient re-organizations to developmental regulation may have led to extinction/irrelevance of fungal lines. The emergence of *brlA* arises as a key event in the re-design of genetic networks and the ability to generate asexual CM structures in eurotiales. It is tempting to suggest that the promoter of *brlA* and its gene products developed, respectively, the ability to recruit upstream and transcriptionally control downstream TFs. Processes such as this could also redirect the role of proteins that are widely conserved in fungi (such as the psi factor-producing oxygenase PpoA or the oxidoreductase GmcA) or conserved in higher eukaryotes (such as HymA). Consequently, we firmly believe that the elucidation of the mechanisms enabling the transition to asexual development in fungal species such as *A. nidulans* could make an important contribution to the understanding of the evolution of CM.

## Acknowledgements

Work at the UPV/EHU lab was funded by the University of the Basque Country (grant EHUA15/08). Work at CIB-CSIC was funded by MINECO/FEDER/EU (grant BFU2015-66806-R).

---

**<u>Figure S1:</u> Sequence analysis of MoConX2.** A) Annotation of the mgg_02775 transcript, coding for MoConX2, and the position of introns, confirmed by RNAseq. Taken from the FungiDB database. B) BLAST results for MoConX2, taken from the NCBI BLAST algorithm website (https://blast.ncbi.nlm.nih.gov/Blast.cgi?PAGE=Proteins). Only the best hits are shown. C) Alignment (Genedoc software) of MoConX2 and the best hits of panel B. The position of the zinc finger-type transcriptional regulatory domain is indicated by a red line. D) Phylogenetic tree (Mega software) showing the sequence divergence of MoConX2 compared to the best hits in panel B.

## References.

Adams TH, Boylan MT, Timberlake WE. 1988. *brlA* is necessary and sufficient to direct conidiophore development in *Aspergillus nidulans*. Cell. 54:353–362.

Adams TH, Wieser JK, Yu J-H. 1998. Asexual sporulation in *Aspergillus nidulans*. Microbiol Mol Biol Rev. 62:35–54.

Aguirre J. 1993. Spatial and temporal controls of the *Aspergillus brlA* developmental regulatory gene. Mol Microbiol. 8:211–218.

Ahmed YL, Gerke J, Park H-S, Bayram Ö, Neumann P, Ni M, Dickmanns A, Kim SC, Yu J-H, Braus GH, Ficner R. 2013. The Velvet family of fungal regulators contains a DNA-binding domain structurally similar to NF-κB. PLoS Biol. 11:e1001750.

Alkhayyat F, Ni M, Kim SC, Yu J-H. 2015. The WOPR domain protein OsaA orchestrates development in *Aspergillus nidulans*. PLoS One. 10:e0137554.

Andrianopoulos A. 1991. ATTS, a new and conserved DNA binding domain. PLANT CELL ONLINE. 3:747–748.

Andrianopoulos A, Timberlake WE. 1994. The *Aspergillus nidulans abaA* gene encodes a transcriptional activator that acts as a genetic switch to control development. Mol Cell Biol. 14:2503–2515.

Antal Z, Rascle C, Cimerman A, Viaud M, Billon-Grand G, Choquer M, Bruel C. 2012. The Homeobox BcHOX8 gene in *Botrytis Cinerea* regulates vegetative growth and morphology. PLoS One. 7:e48134.

Bar-Yam Y, Harmon D, de Bivort B. 2009. Attractors and Democratic Dynamics. Science (80-). 323:1016 LP–1017.

Bayram Ö, Bayram ÖS, Ahmed YL, Maruyama J, Valerius O, Rizzoli SO, Ficner R, Irniger S, Braus GH. 2012. The *Aspergillus nidulans* MAPK module AnSte11-Ste50-Ste7-Fus3 controls development and secondary metabolism. PLoS Genet. 8:e1002816.

Bayram Ö, Braus GH. 2012. Coordination of secondary metabolism and development in fungi: the velvet family of regulatory proteins. FEMS Microbiol Rev. 36:1–24.

Bayram Ö, Braus GH, Fischer R, Rodriguez-Romero J. 2010. Spotlight on *Aspergillus nidulans* photosensory systems. Fungal Genet Biol. 47:900–908.

Bayram Ö, Krappmann S, Ni M, Bok JW, Helmstaedt K, Valerius O, Braus-Stromeyer S, Kwon N-J, Keller NP, Yu J-H, Braus GH. 2008. VelB/VeA/LaeA complex coordinates light signal with fungal development and secondary metabolism. Science (80-). 320:1504 LP–1506.

Bebber DP, Holmes T, Gurr SJ. 2014. The global spread of crop pests and pathogens. Glob Ecol Biogeogr. 23:1398–1407.

Bhattacharya S. 2017. Deadly new wheat disease threatens Europe’s crops. Nature. 542:145–146.

Boni AC, Ambrósio DL, Cupertino FB, Montenegro-Montero A, Virgilio S, Freitas FZ, Corrocher FA, Gonçalves RD, Yang A, Weirauch MT, et al. 2018. *Neurospora crassa* developmental control mediated by the FLB-3 transcription factor. Fungal Biol. 122:570–582.

Boylan MT, Mirabito PM, Willett CE, Zimmerman CR, Timberlake WE. 1987. Isolation and physical characterization of three essential conidiation genes from *Aspergillus nidulans*. Mol Cell Biol. 7:3113–3118.

Brandhoff B, Simon A, Dornieden A, Schumacher J. 2017. Regulation of conidiation in *Botrytis cinerea* involves the light-responsive transcriptional regulators BcLTF3 and BcREG1. Curr Genet. 63:931–949.

Busby TM, Miller KY, Miller BL. 1996. Suppression and enhancement of the *Aspergillus nidulans medusa* mutation by altered dosage of the *bristle* and *stunted* Genes. Genetics. 143:155 LP–163.

Callaway E. 2016. Devastating wheat fungus appears in Asia for first time. Nature. 532:421–422.

Cánovas D, Marcos AT, Gacek A, Ramos MS, Gutiérrez G, Reyes-Domínguez Y, Strauss J. 2014. The histone acetyltransferase GcnE (GCN5) plays a central role in the regulation of *Aspergillus* asexual development. Genetics. 197:1175–89.

Cao H, Huang P, Zhang L, Shi Y, Sun D, Yan Y, Liu X, Dong B, Chen G, Snyder JH, et al. 2016. Characterization of 47 Cys2□His2 zinc finger proteins required for the development and pathogenicity of the rice blast fungus *Magnaporthe oryzae*. New Phytol. 211:1035–1051.

de Castro F, Bolker BM. 2005. Parasite establishment and host extinction in model communities. Oikos. 111:501–513.

Cohrs KC, Simon A, Viaud M, Schumacher J. 2016. Light governs asexual differentiation in the grey mould fungus *Botrytis cinerea* via the putative transcription factor BcLTF2. Environ Microbiol. 18:4068–4086.

Cortese MS, Etxebeste O, Garzia A, Espeso EA, Ugalde U. 2011. Elucidation of functional markers from *Aspergillus nidulans* developmental regulator FlbB and their phylogenetic distribution. PLoS One. 6:e17505.

Editorial Nature Microbiology. 2017. Stop neglecting fungi. Nat Microbiol. 2; 17120.

Etxebeste O, Garzia A, Espeso EA, Ugalde U. 2010. *Aspergillus nidulans* asexual development: Making the most of cellular modules. Trends Microbiol. 18:569–576.

Etxebeste O, Herrero-García E, Cortese MS, Garzia A, Oiartzabal-Arano E, de los Ríos V, Ugalde U, Espeso EA. 2012. GmcA is a putative glucose-methanol-choline oxidoreductase required for the induction of asexual development in *Aspergillus nidulans*. PLoS One. 7:e40292.

Etxebeste O, Ni M, Garzia A, Kwon NJ, Fischer R, Yu JH, Espeso EA, Ugalde U. 2008. Basic-zipper-type transcription factor FlbB controls asexual development in *Aspergillus nidulans*. Eukaryot Cell. 7:38–48.

Fischer R, Kües U. 2006. Asexual sporulation in mycelial fungi. In: Kües U, Fischer R, editors. Growth, Differ Sex. Berlin, Heidelberg: Springer Berlin Heidelberg; p. 263– 292.

Fisher MC, Henk DA, Briggs CJ, Brownstein JS, Madoff LC, McCraw SL, Gurr SJ. 2012. Emerging fungal threats to animal, plant and ecosystem health. Nature. 484:186– 194.

Garzia A, Etxebeste O, Herrero-Garcia E, Fischer R, Espeso EA, Ugalde U. 2009. *Aspergillus nidulans* FlbE is an upstream developmental activator of conidiation functionally associated with the putative transcription factor FlbB. Mol Microbiol. 71:172–184.

Garzia A, Etxebeste O, Herrero-García E, Ugalde U, Espeso EA. 2010. The concerted action of bZip and cMyb transcription factors FlbB and FlbD induces *brlA* expression and asexual development in *Aspergillus nidulans*. Mol Microbiol. 75:1314–1324.

Garzia A, Etxebeste O, Rodríguez-Romero J, Fischer R, Espeso EA, Ugalde U. 2013. Transcriptional changes in the transition from vegetative cells to asexual development in the model fungus *Aspergillus nidulans*. Eukaryot Cell. 12:311–321.

Gonçalves AP, Heller J, Daskalov A, Videira A, Glass NL. 2017. Regulated forms of cell death in fungi. Front Microbiol. 8:1837.

Han K, Han K, Yu J, Chae K, Jahng K, Han D. 2001. The *nsdD* gene encodes a putative GATA-type transcription factor necessary for sexual development of *Aspergillus nidulans*. Mol Microbiol. 41:299–309.

Han S, Navarro J, Greve RA, Adams TH. 1993. Translational repression of *brlA* expression prevents premature development in *Aspergillus*. EMBO J. 12:2449–2457.

Hawksworth DL, Lücking R. 2017. Fungal diversity revisited: 2.2 to 3.8 million species. Microbiol Spectr. 5.

Herrero-Garcia E, Perez-de-Nanclares-Arregi E, Cortese MS, Markina-Iñarrairaegui A, Oiartzabal-Arano E, Etxebeste O, Ugalde U, Espeso EA. 2015. Tip-to-nucleus migration dynamics of the asexual development regulator FlbB in vegetative cells. Mol Microbiol. 98:607–24.

Huerta-Cepas J, Forslund K, Coelho LP, Szklarczyk D, Jensen LJ, von Mering C, Bork P. 2017. Fast genome-wide functional annotation through orthology assignment by eggNOG-Mapper. Mol Biol Evol. 34:2115–2122.

Huerta-Cepas J, Szklarczyk D, Forslund K, Cook H, Heller D, Walter MC, Rattei T, Mende DR, Sunagawa S, Kuhn M, et al. 2016. eggNOG 4.5: a hierarchical orthology framework with improved functional annotations for eukaryotic, prokaryotic and viral sequences. Nucleic Acids Res. 44:D286–D293.

Huerta-Cepas J, Szklarczyk D, Heller D, Hernández-Plaza A, Forslund SK, Cook H, Mende DR, Letunic I, Rattei T, Jensen LJ, et al. 2019. eggNOG 5.0: a hierarchical, functionally and phylogenetically annotated orthology resource based on 5090 organisms and 2502 viruses. Nucleic Acids Res. 47:D309–D314.

Islam TM, Croll D, Gladieux P, Soanes DM, Persoons A, Bhattacharjee P, Shaid Hossain M, Rani Gupta D, Mahbubur Rahman M, Golam Mahboob M, et al. 2016. Emergence of wheat blast in Bangladesh was caused by a South American lineage of *Magnaporthe oryzae*. BMC Biol. 14.

Johnson AJ. 2016. Artisanal food microbiology. Nat Microbiol. 1:16039.

Jothi R, Balaji S, Wuster A, Grochow JA, Gsponer J, Przytycka TM, Aravind L, Babu MM. 2009. Genomic analysis reveals a tight link between transcription factor dynamics and regulatory network architecture. Mol Syst Biol. 5:294.

Karos M, Fischer R. 1996. hymA (hypha-like metulae), a new developmental mutant of Aspergillus nidulans. Microbiology. 142:3211–3218.

Kim H-R, Chae K-S, Han K-H, Han D-M. 2009. The *nsdC* gene encoding a putative C2H2-type transcription factor is a key activator of sexual development in *Aspergillus nidulans*. Genetics. 182:771 LP–783.

Kirk P, Cannon P, Minter D, Stalpers J. 2008. Dictionary of the Fungi (10th Edition). 10th ed. Wallingford: CAN International.

Krizsán K, Almási É, Merényi Z, Sahu N, Virágh M, Kószó T, Mondo S, Kiss B, Bálint B, Kües U, et al. 2019. Transcriptomic atlas of mushroom development reveals conserved genes behind complex multicellularity in fungi. Proc Natl Acad Sci.:201817822.

Kües U, Badalyan SM, Gießler A, Dörnte B. 2016. Asexual sporulation in Agaricomycetes. In: Wendland J, editor. Growth, Differ Sex. Cham: Springer International Publishing; p. 269–328.

Kwon N-J, Garzia A, Espeso EA, Ugalde U, Yu J-H. 2010. FlbC is a putative nuclear C2H2 transcription factor regulating development in *Aspergillus nidulans*. Mol Microbiol. 77:1203–1219.

Lee BN, Adams TH. 1994. Overexpression of *flbA*, an early regulator of *Aspergillus* asexual sporulation, leads to activation of *brlA* and premature initiation of development. Mol Microbiol. 14:323–334.

Lee M-K, Kwon N-J, Choi JM, Lee I-S, Jung S, Yu J-H. 2014. NsdD is a key repressor of asexual development in *Aspergillus nidulans*. Genetics. 197:159–173.

Lee M-K, Kwon N-J, Lee I-S, Jung S, Kim S-C, Yu J-H. 2016. Negative regulation and developmental competence in *Aspergillus*. Sci Rep. 6:28874.

Li F, Shi H-Q, Ying S-H, Feng M-G. 2015. WetA and VosA are distinct regulators of conidiation capacity, conidial quality, and biological control potential of a fungal insect pathogen. Appl Microbiol Biotechnol. 99:10069–10081.

Lind AL, Lim FY, Soukup AA, Keller NP, Rokas A. 2018. An LaeA- and BrlA-dependent cellular network governs tissue-specific secondary metabolism in the human pathogen Aspergillus fumigatus. mSphere. 3:e00050–18.

Lockhart SR, Etienne KA, Vallabhaneni S, Farooqi J, Chowdhary A, Govender NP, Colombo AL, Calvo B, Cuomo CA, Desjardins CA, et al. 2017. Simultaneous emergence of multidrug-resistant *Candida auris* on 3 continents confirmed by whole-genome sequencing and epidemiological analyses. Clin Infect Dis. 64:134–140.

Mah J-H, Yu J-H. 2006. Upstream and downstream regulation of asexual development in *Aspergillus fumigatus*. Eukaryot Cell. 5:1585–1595.

Malaker PK, Barma NCD, Tiwari TP, Collis WJ, Duveiller E, Singh PK, Joshi AK, Singh RP, Braun H-J, Peterson GL, et al. 2016. First report of wheat blast caused by *Magnaporthe oryzae* pathotype *triticum* in Bangladesh. Plant Dis. 100:2330.

Marshall MA, Timberlake WE. 1991. *Aspergillus nidulans wetA* activates spore-specific gene expression. Mol Cell Biol. 11:55–62.

McCallum H, Dobson A. 1995. Detecting disease and parasite threats to endangered species and ecosystems. Trends Ecol Evol. 10:190–194.

Meyer V, Andersen MR, Brakhage AA, Braus GH, Caddick MX, Cairns TC, de Vries RP, Haarmann T, Hansen K, Hertz-Fowler C, et al. 2016. Current challenges of research on filamentous fungi in relation to human welfare and a sustainable bio-economy: a white paper. Fungal Biol Biotechnol. 3:6.

Miller KY, Wu J, Miller BL. 1992. StuA is required for cell pattern formation in *Aspergillus*. Genes Dev. 6:1770–1782.

Mims CW, Richardson EA, Timberlake WE. 1988. Ultrastructural analysis of conidiophore development in the fungus *Aspergillus nidulans* using freeze-substitution. Protoplasma. 144:132–141.

Mooney JL, Yager LN. 1990. Light is required for conidiation in *Aspergillus nidulans*. Genes Dev. 4:1473–1482.

Nagy LG. 2017. Evolution: Complex multicellular life with 5,500 genes. Curr Biol. 27:R609–R612.

Nagy LG, Gábor M. K, Krisztina K. 2018. Complex multicellularity in fungi: evolutionary convergence, single origin, or both? Biol Rev. 93:1778–1794.

Nguyen TA, Cissé OH, Yun Wong J, Zheng P, Hewitt D, Nowrousian M, Stajich JE, Jedd G. 2017. Innovation and constraint leading to complex multicellularity in the Ascomycota. Nat Commun. 8:14444.

Ni M, Yu J-H. 2007. A novel regulator couples sporogenesis and trehalose biogenesis in *Aspergillus nidulans*. PLoS One. 2:e970.

Oiartzabal-Arano E, Garzia A, Gorostidi A, Ugalde U, Espeso EA, Etxebeste O. 2015. Beyond asexual development: Modifications in the gene expression profile caused by the absence of the *Aspergillus nidulans* transcription factor FlbB. Genetics. 199:1127– 42.

Oiartzabal-Arano E, Perez-de-Nanclares-Arregi E, Espeso EA, Etxebeste O. 2016. Apical control of conidiation in *Aspergillus nidulans*. Curr Genet. 62:371–377.

Ojeda-López M, Chen W, Eagle CE, Gutiérrez G, Jia WL, Swilaiman SS, Huang Z, Park H-S, Yu J-H, Cánovas D, Dyer PS. 2018. Evolution of asexual and sexual reproduction in the aspergilli. Stud Mycol. 91:37–59.

Park H-S, Nam T-Y, Han K-H, Kim SC, Yu J-H. 2014. VelC positively controls sexual development in *Aspergillus nidulans*. PLoS One. 9:e89883.

Park H-S, Yu J-H. 2012. Genetic control of asexual sporulation in filamentous fungi. Curr Opin Microbiol. 15:669–677.

Pócsi I, Leiter E, Kwon N-J, Shin K-S, Kwon G-S, Pusztahelyi T, Emri T, Abuknesha RA, Price RG, Yu J-H. 2009. Asexual sporulation signalling regulates autolysis of *Aspergillus nidulans* via modulating the chitinase ChiB production. J Appl Microbiol. 107:514–523.

Purschwitz J, Müller S, Kastner C, Schöser M, Haas H, Espeso EA, Atoui A, Calvo AM, Fischer R. 2008. Functional and physical interaction of blue- and red-light sensors in *Aspergillus nidulans*. Curr Biol. 18:255–259.

Rodriguez-Romero J, Hedtke M, Kastner C, Müller S, Fischer R. 2010. Fungi, hidden in soil or up in the air: light makes a difference. Annu Rev Microbiol. 64:585–610.

Schumacher J. 2017. How light affects the life of *Botrytis*. Fungal Genet Biol. 106:26– 41.

Selitrennikoff CP, Nelson RE, Siegel RW. 1974. Phase-specific genes for macroconidiation in Neurospora crassa. Genetics. 78:679 LP–690.

Sewall TC, Mims CW, Timberlake WE. 1990. Conidium differentiation in *Aspergillus nidulans* wild-type and wet-white (*wetA*) mutant strains. Dev Biol. 138:499–508.

Son H, Kim M-G, Chae S-K, Lee Y-W. 2014. FgFlbD regulates hyphal differentiation required for sexual and asexual reproduction in the ascomycete fungus *Fusarium graminearum*. J Microbiol. 52:930–939.

Spatafora JW, Aime MC, Grigoriev I V., Martin F, Stajich JE, Blackwell M. 2017. The fungal tree of life: from molecular systematics to genome-scale phylogenies. Microbiol Spectr. 5.

Stajich JE. 2017. Fungal genomes and insights into the evolution of the kingdom. Microbiol Spectr. 5.

Stinnett SM, Espeso EA, Cobeño L, Araújo□Bazán L, Calvo AM. 2006. *Aspergillus nidulans* VeA subcellular localization is dependent on the importin-α carrier and on light. Mol Microbiol. 63:242–255.

Tang W, Ru Y, Hong L, Zhu Q, Zuo R, Guo X, Wang J, Zhang H, Zheng X, Wang P, Zhang Z. 2014. System-wide characterization of bZIP transcription factor proteins involved in infection-related morphogenesis of *Magnaporthe oryzae*. Environ Microbiol. 17:1377–1396.

Tiley AMM, Foster GD, Bailey AM. 2018. Exploring the genetic regulation of asexual sporulation in *Zymoseptoria tritici*. Front Microbiol. 9:1859.

Tsitsigiannis DI, Zarnowski R, Keller NP. 2004. The lipid body protein, PpoA, coordinates sexual and asexual sporulation in *Aspergillus nidulans*. J Biol Chem. 279:11344–11353.

Tüncher A, Reinke H, Martic G, Caruso Maria L, Axe lA. B. 2004. A basic□region helix–loop–helix protein□encoding gene (*devR*) involved in the development of *Aspergillus nidulans*. Mol Microbiol. 52:227–241.

Vallim MA, Miller KY, Miller BL. 2002. *Aspergillus* SteA (Sterile12□like) is a homeodomain□C2/H2□Zn+2 finger transcription factor required for sexual reproduction. Mol Microbiol. 36:290–301.

Vivancos AP, Castillo EA, Jones N, Ayté J, Hidalgo E. 2004. Activation of the redox sensor Pap1 by hydrogen peroxide requires modulation of the intracellular oxidant concentration. Mol Microbiol. 52:1427–1435.

de Vries RP, Riley R, Wiebenga A, Aguilar-Osorio G, Amillis S, Uchima CA, Anderluh G, Asadollahi M, Askin M, Barry K, et al. 2017. Comparative genomics reveals high biological diversity and specific adaptations in the industrially and medically important fungal genus *Aspergillus*. Genome Biol. 18:28.

Wieser J, Lee BN, Fondon JW, Adams TH. 1994. Genetic requirements for initiating asexual development in *Aspergillus nidulans*. Curr Genet. 27:62–69.

Wu M-Y, Mead ME, Lee M-K, Ostrem Loss EM, Kim S-C, Rokas A, Yu J-H. 2018. Systematic dissection of the evolutionarily conserved WetA developmental regulator across a genus of filamentous fungi. MBio. 9:e01130–18.

Yu JH, Wieser J, Adams TH. 1996. The *Aspergillus* FlbA RGS domain protein antagonizes G protein signaling to block proliferation and allow development. EMBO J. 15:5184–5190.

