## Supplementary figures and images for "Rewiring of transcriptional networks as a major event leading to the diversity of asexual multicellularity in fungi"

### Figure S1

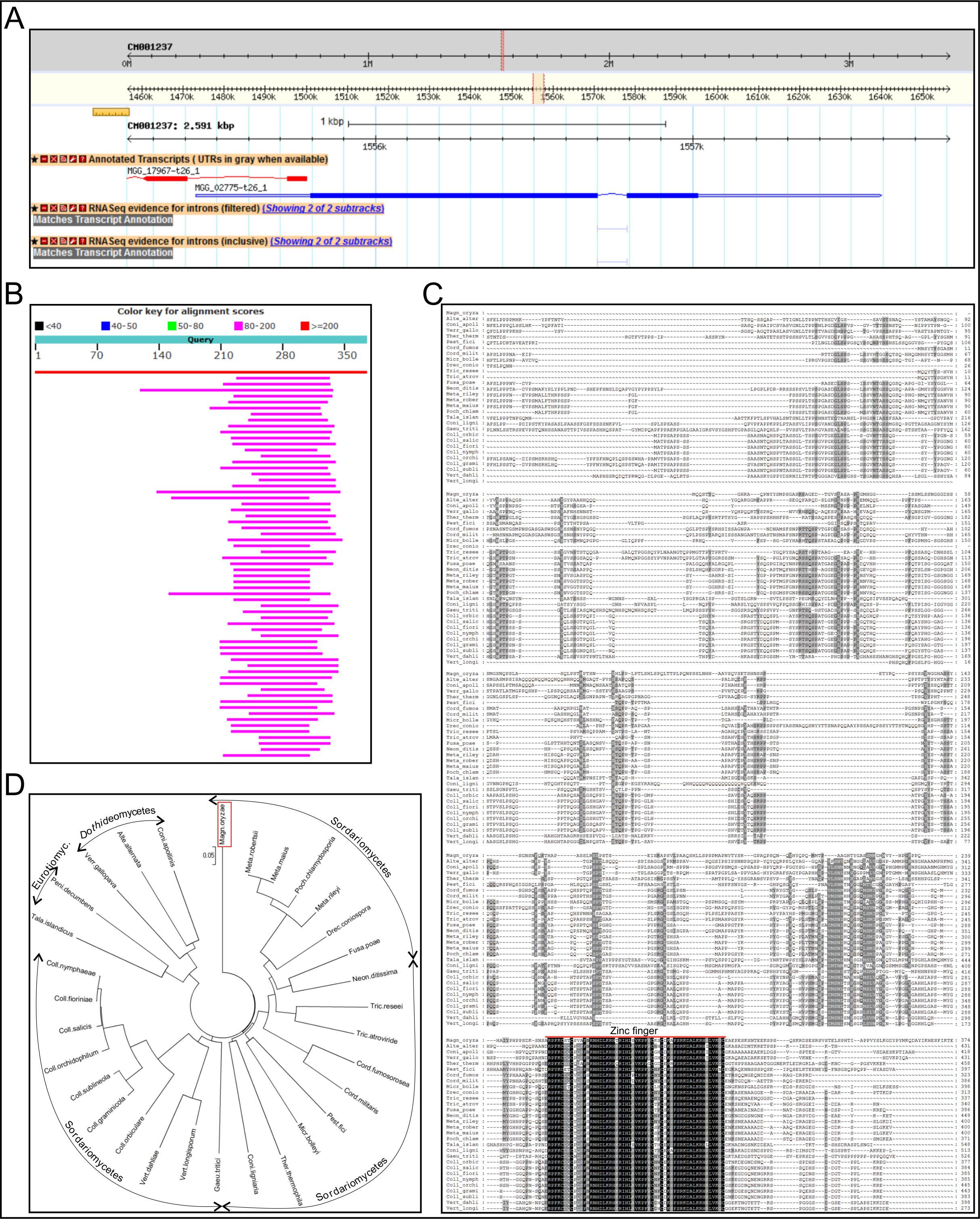
